# A popular Indian clove-based mosquito repellent is a false economy

**DOI:** 10.1101/689836

**Authors:** Kaiming Tan, Gabriel B. Faierstein, Pingxi Xu, Rosângela M.R. Barbosa, Garrison K. Buss, Walter S. Leal

**Author notes:** Corresponding author: Walter S. Leal, Department of Molecular and Cellular Biology, University of California-Davis, Davis CA 95616 USA. Department of Molecular and Cellular Physiology, Stanford University, Stanford, CA 94305, U.S.A.

## Abstract

Insect repellents are important prophylactic tools for travelers to and populations living in endemic areas of malaria, dengue, encephalitis, and other vector-borne diseases, and the first line of defense against emerging arboviruses. However, the cost of daily applications of even the most affordable and the gold standard of insect repellents, DEET, is still high for low-income populations where repellents are needed the most. An Indian clove-based homemade recipe has been presented as a panacea. We analyzed this homemade repellent and confirmed by behavioral measurements and odorant receptor responses that eugenol is the active ingredient in this formulation. Prepared as advertised, this homemade repellent is ineffective, whereas 5X more concentrated extracts from the brand most enriched in eugenol showed moderate repellency activity against *Culex quinquefasciatus* and *Aedes aegypti*. DEET showed higher performance when compared to the 5X concentrated formulation and is available in the same market at a lower price than the cost of the ingredients to prepare the homemade formulation.

## Introduction

Diseases transmitted by mosquitoes destroy more lives on a year basis than war, gun violence, and other human maladies combined^1^. On the top of the list of the most devastating diseases transmitted by mosquitoes is malaria, which is caused by parasites that are transmitted to people through the bites of infected female *Anopheles* mosquitoes and led to an estimated 435,000 deaths with 219 million cases in 87 countries in 2017 alone^2^. A close second is dengue, which is caused by a virus primarily transmitted by the yellow fever mosquito, *Aedes (=Stegomyia) aegypti*. Severe dengue is a leading cause of serious illness and death among children in some Asian and Latin American countries. Dengue is now endemic in more than 100 countries and about half of the world population is at risk^3^. Zika became notorious not only because of its explosive outbreak in Latin America, but also for causing a congenital Zika syndrome. There are many other mosquito-borne diseases, including chikungunya, West Nile, yellow fever, and Mayaro just to cite a few. The Southern house mosquito, *Culex quinquefasciatus*, transmits the nematode *Wuchereria bancrofti* that causes the debilitating lymphatic filariasis – a neglected tropical disease. Viruses transmitted by *Culex* mosquitoes are West Nile (WNV), St. Louis encephalitis (SLEv), Western equine encephalitis (WEEv), and Rift Valley fever virus.

The first line of defense against infected mosquitoes by people living in or traveling to endemic areas is to use insect repellents. With the growing number of emerging mosquito-borne diseases, it is not surprising that consumption of repellents is in an upward trend. In the United States alone consumption was 16.7% higher in 2018 than in 2014. It is projected that more than 200 million Americans (equivalent to 60% of the population) will use insect repellents in 2020 ^4^. There are a number of commercially available mosquito repellents and homemade products, with the synthetic repellent DEET (N,N,-diethyl-3-methylbenzamide) being the gold standard ^1^. DEET works primarily as a spatial ^5,6^ and a contact repellent ^7,8^, but it is also a feeding deterrent ^9^. Mosquitoes attracted to humans may not land on DEET-treated skin because they smell the repellent at very short distances and steer away, but if this spatial repellency fails, mosquitoes are repelled when landing on DEET-treated skin. Albeit effective, DEET is not widely used in endemic areas because it may not be affordable to the needy populations. Also, among those who can afford DEET there are people averse to synthetic products. Therefore, there is a growing number of homemade and natural product-based mosquito repellents aimed at those who cannot afford DEET and those who do not like DEET, respectively. In Brazil, for example, one of the most popular homemade repellent is based on alcoholic extract of Indian clove (Cravo-da-Índia in Portuguese). Here, we compared the repellency of this homemade repellent with that of DEET and report that the homemade recipe is more costly and less effective than DEET.

## Results and Discussion

### Comparing ethanolic extracts of three commercially available Indian clover buds

There are various recipes of Indian clove-based repellents. In general, they call for an extraction of whole clover buds by soaking 10 or 30 g in 500 ml of ethanol for 4 days (shaking twice a day) and then filtering and mixing the supernatant with baby oil. Indian clove is readily available in supermarkets in Brazil where they are used primarily for culinary purposes. We extracted whole clover buds from three different commercial brands in Brazil, namely Portuense, Beija Flor, and Kitano following the popular protocol and using the higher dose (30 g per 500 ml of ethanol). Then, we compared their gas chromatographic profiles (Fig. 1). Compounds were identified by their mass spectra (Fig. 2) and retention times using authentic standards. The major constituents of the ethanolic extracts in all these samples were eugenol, followed by ß-caryophyllene, and eugenyl acetate, in case of Portuense and Beija Flor and eugenyl acetate and ß-caryophyllene in case of Kitano (Fig. 1,2). Two other minor constituents were α-humulene and caryophyllene oxide (Fig. 1,2). This observed variation in composition is consistent with earlier analysis of clove oil, ^10–13^ although it is worth noting that these ethanolic extracts and essential oils (generated by steam distillation) are quite different. It has been previously reported that eugenol, isolated from an essential oil of *Monarda bradburiana*, is a mosquito repellent ^14^. As far as the percent composition of eugenol is concerned, the Indian clove buds we analyzed appeared in the order: Kitano > Portuense > Beja Flor (Fig. 1). However, all these extracts had very low concentrations of eugenol thus suggesting that following the original protocol (even with 30 g per 500 ml) may lead to inactive extracts. Using an arm-in-cage set-up, Miot and collaborators found no significant difference in repellency when comparing arms treated with this homemade recipe and untreated arms ^15^. By contrast Affonso and collaborators reported that clove buds ethanolic extracts obtained by static maceration showed repellence activity against the yellow fever mosquito, *Ae. aegypti* ^16^. It is likely that both reports are accurate, the difference being the method of extraction. We, therefore, changed the original protocol to obtain 5X more concentrated samples of Indian clove buds, ie, 30 g per 100 ml. This could be done by increasing the amount of buds or by reducing the volume of ethanol. These 5X extracts were obtained with enough solvent to cover the buds, but following the same protocol, ie, shaking twice a day, but no maceration. We compared the three 5X formulations in a time course of 4 (standard time), 5, and 6 days. We aimed at obtaining extracts with at least 10 mg of eugenol per ml, which is the minimal desired concentration (1%). In our surface landing and feeding assays, a dose of 1% is equivalent to approximately 6.3% dose when tested in the arm-in-cage assays ^17^, which in turn is the most common dose of DEET-based repellents in the market. ^17^

**Figure 1.**
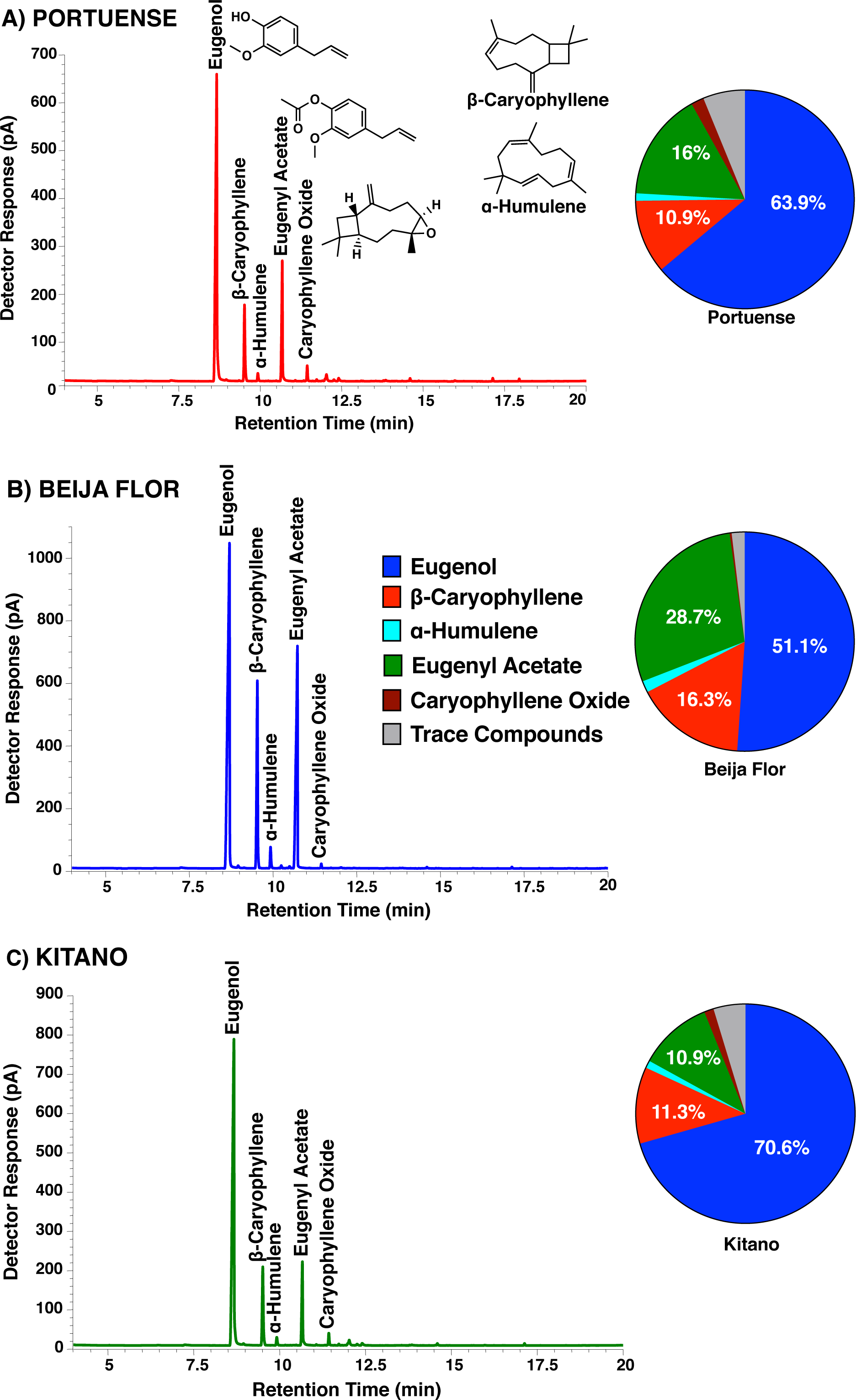
Gas chromatographic profiles obtained with three samples of Indian clove buds from supermarkets in Brazil. (A) Portuense, (B) Beija Flor, and (C) Kitano. Each peak was identified on the basis of its mass spectrum and retention time by comparison with authentic standards. The solvent peak was omitted for clarity. The compositions of the major constituents in these 1X extracts are displayed in pie graphics.

**Figure 2.**
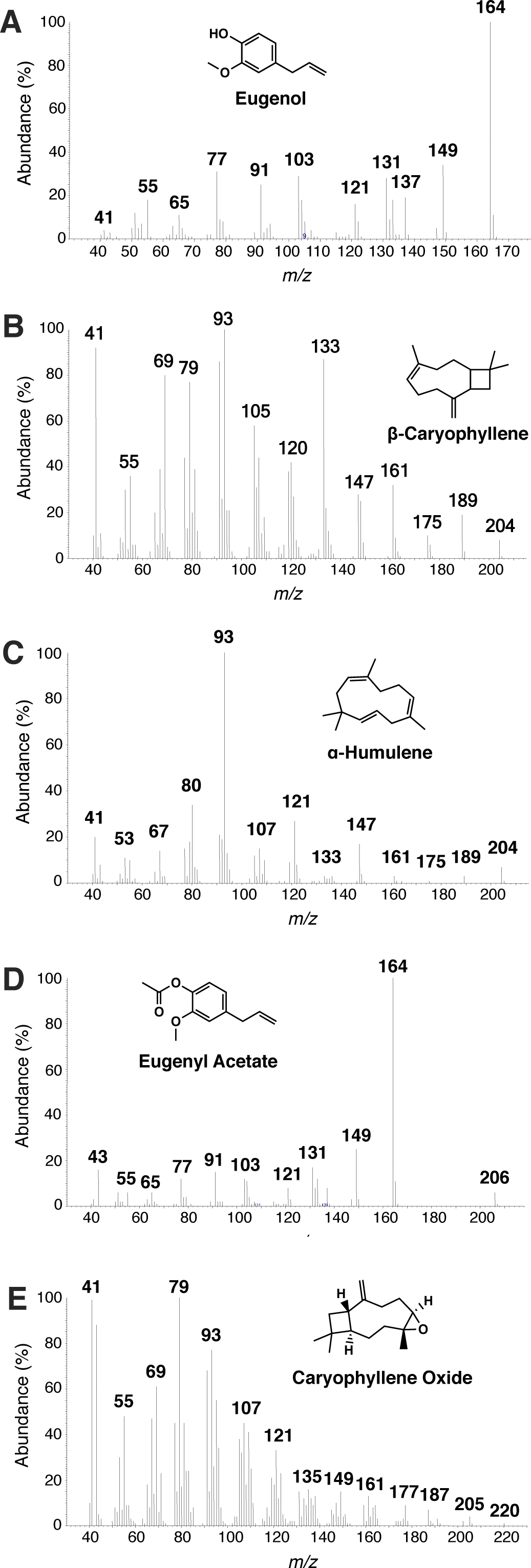
Mass spectra of the major constituents of Indian clove bud extracts. (A) Eugenol, (B) β-caryophyllene, (C) α-humulene, (D) eugenyl acetate, and (E) caryophyllene oxide. The respective mass spectra obtained with authentic standards were undistinguishable from MS of the natural products.

Our analysis (Fig. 3) showed a time-dependent increase in the concentration of eugenol in the extracts from all brands. Levels of eugenol in Portuense were very low even in 6-day extracts, whereas Beija Flor reached the minimal desirable level within 4 days; this brand produced the highest levels of eugenol throughout the tested period (Fig. 3). Kitano gave intermediate results. We then focused on Beija Flor 5X recipe, 4-day extracts for all behavioral studies.

**Figure 3.**
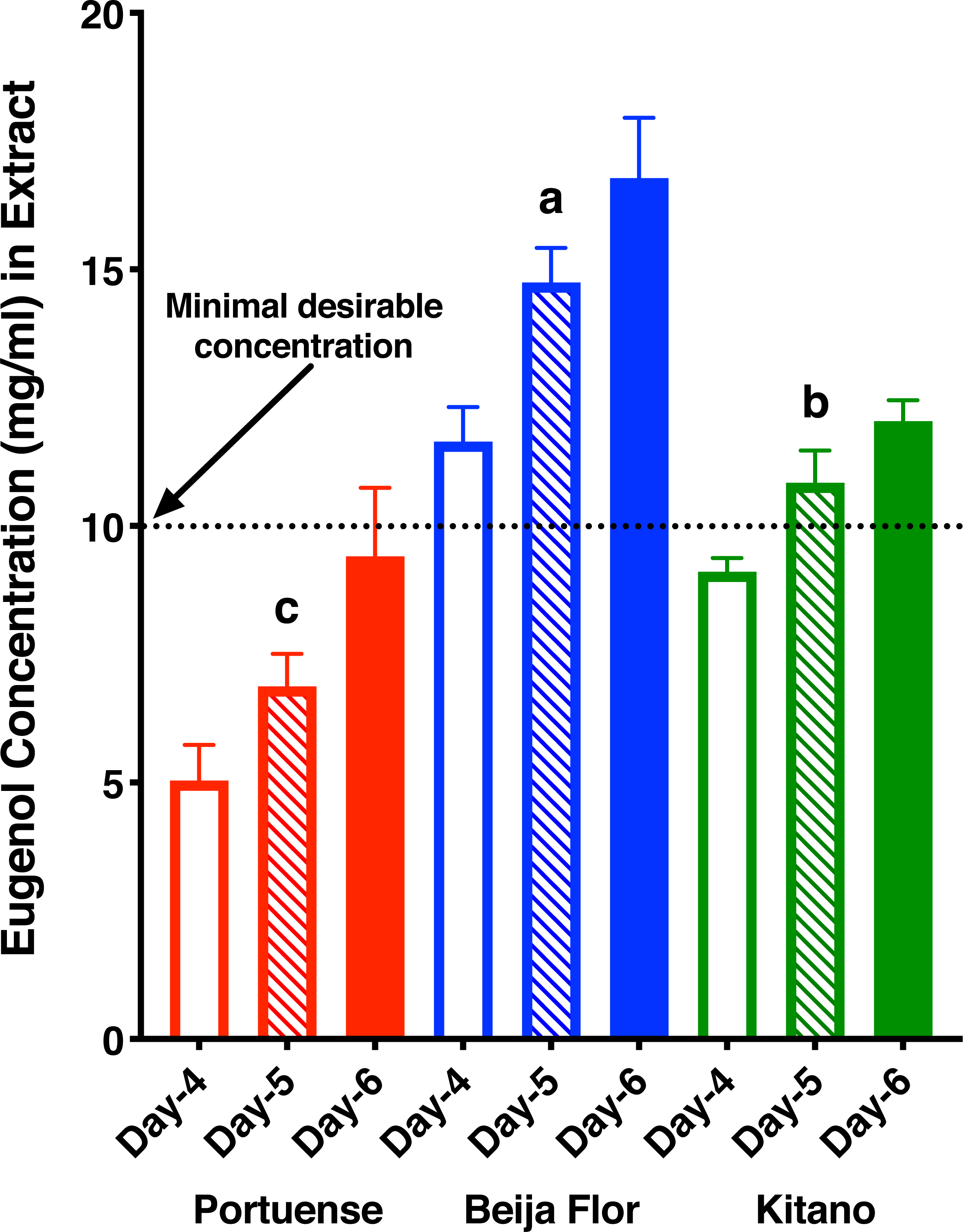
Time course analysis of 5X homemade recipes. Minimal desirable concentration (10 mg/ml, 1%) is a typical concentration that natural products, when active, shows repellency activity. In our assays this concentration is equivalent to 6-7% in arm-in cage repellent assays. The latter is typically found in DEET-based commercial repellents. Amounts of eugenol (Mean ± SEM) in triplicate extracts from 4, 5 and 6 days. Levels of eugenol in Day-5 samples were compared by Brown-Forsythe (F* = 37.17, P = 0.0004) and Welch (W = 31.01, P = 0.037) ANOVA. Beija Flor vs. Portuense, adjusted P = 0.0027; Portuense vs. Kitano, adjusted P = 0.0284; Beija Flor vs. Kitano, P = 0.0334). Concentrations at Day-6 were higher, but not significantly different from their respective Day-5 samples (Portuense, P = 0.2178; Beija Flor, P = 0.2966; Kitano, P = 0.2352).

### *Cx. quinquefasciatus* responses to an improved homemade formulation and its constituents

First, we compared in our surface-landing and feeding assay ^6,17^, repellency elicited by 5X Beija Flor extracts against the Southern house mosquito, *Culex quinquefasciatus*. With 5X extracts, in average (*n* = 9) 19.6±2.7 female mosquitoes per trial landed on the control (solvent only) side of the arena, whereas 5.75±0.81 mosquitoes per trial landed on the clover side of the extract. Under similar conditions, only 2±0.41 mosquitoes per trial landed on the DEET 1% side of the arena, whereas 18.75±2.03 females per trial landed on the control side. These repellency activities were then expressed in protection (P = 1-T/C), as recommended by the World Health Organization (WHO) ^18^ and the Environmental Protection Agency (EPA) ^19^. The data show a significantly higher protection elicited by DEET than the 5X homemade repellent (*n* = 9, P = 0.0159, Mann-Whitney test) (Fig. 4A). Next, we tested whether in addition to eugenol the other two major constituents, β-caryophyllene and eugenyl acetate would contribute to the repellency activity of the homemade repellent. There was no significant difference between the two sides of the arena when 1% caryophyllene was tested: control, 7±2.3 females/trial; treatment, 5.25±2.39 females/trial (*n* = 4, P = 0.0.6302, two-tailed, paired t-test). At the same time and with the same group of mosquitoes 1.5±0.5 females/trial landed on the side of the arena treated with DEET 1%; control, 12.25±1.44 females/trial (*n* = 4, P = 0.0061, two-tailed, paired t-test). Likewise, no repellency activity was observed with eugenyl acetate at 1%: control, 10.3±2.8 female mosquitoes per trial; treatment, 6±1.6 female mosquitoes per trial (*n* = 6, P = 0.1875, Wilcoxon matched-pairs signed ranks). Under the same conditions, 22.5±2.16 female mosquitoes per trial landed on the control side of the arena, whereas 1.5±0.6 female mosquitoes per trial landed on DEET 1% side (*n* = 6, P = 0.0312, Wilcoxon matched-pairs signed ranks). By contrast, 1% eugenol showed moderate repellence, but significantly lower protection than 1% DEET (Fig. 4B, *n* = 4, P = 0.0126, Mann-Whitney test).

**Figure 4.**
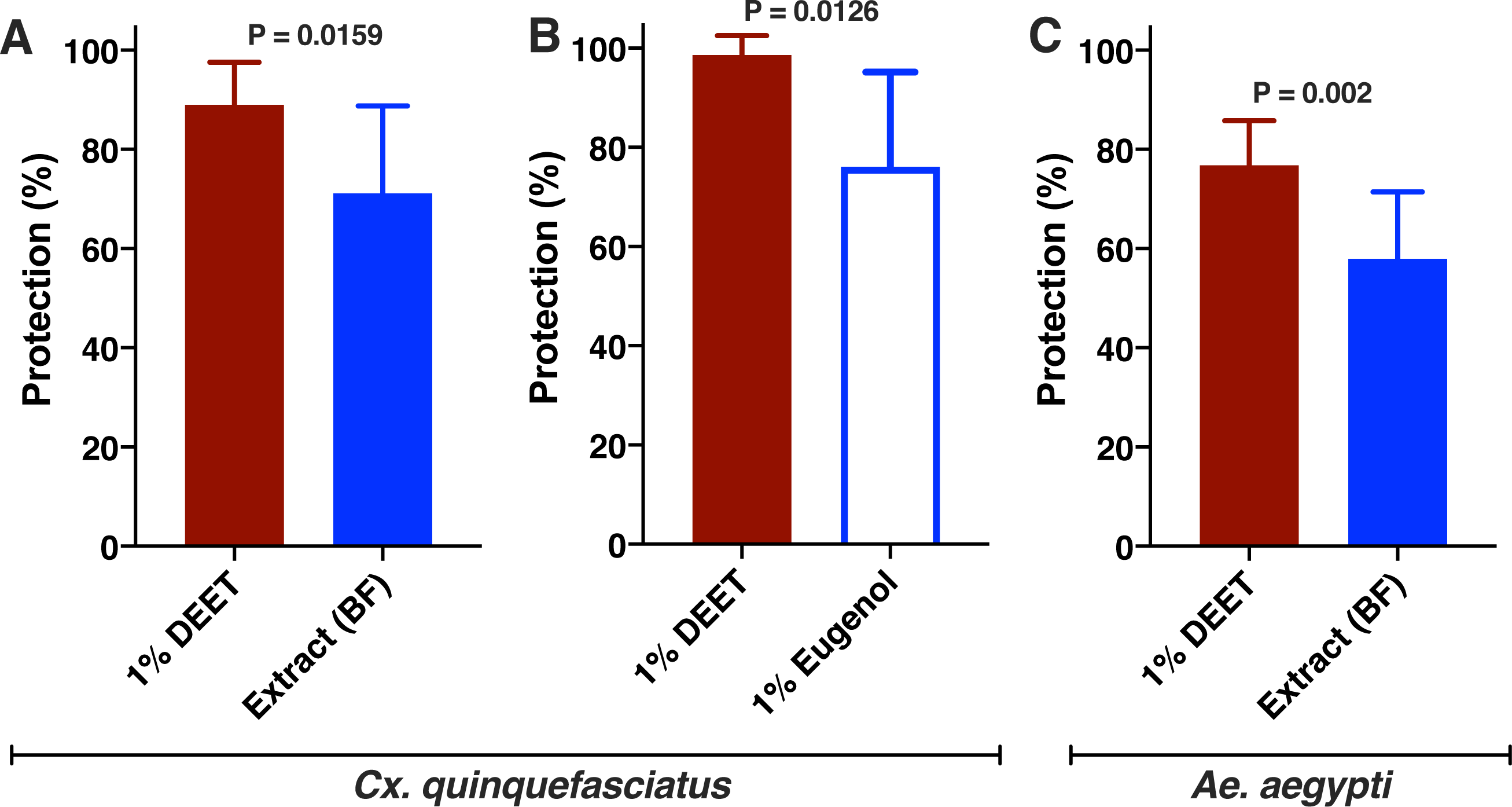
Repellency activity elicited by Indian clove 5X extracts, DEET and eugenol. Data were transformed into protection per WHO^18^ and EPA^19^ recommendations. (A) Comparison of the responses by *Cx. quinquefasciatus* to Beija Flor 5X extracts vs. 1% DEET. (n = 4 for each treatment, P = 0.0159, Mann-Whitney test) and (B) eugenol vs. DEET, both at 1% (n = 8 for each treatment, P = 0.0126, Mann-Whitney test). (C) Comparison of the responses of *Ae. aegypti* to Beija Flor 5X extracts vs. 1% DEET (n = 10, P = 0.002, Mann-Whitney test).

### Odorant receptor sensitive to the active ingredient in Indian clover extracts

Previously we have identified an eugenol-detecting OR in the antennae of *Cx. quinquefasciatus*, CquiOR73, which is narrowly tuned to phenolic compounds^20^. We expressed this receptor, along with the obligatory co-receptor Orco, CquiOrco, in the *Xenopus* oocyte recording system and then compared 5X extracts with known doses of eugenol. We diluted the extracts with Ringer buffer in the ratios 1:100,000, 1:10,000, and 1,1,000. CquiOR73/CquiOrco-expressing oocytes responded to both homemade extract and eugenol in a dose-dependent manner, with saturation at 1mM eugenol (Fig. 5). We observed that a 1:1,000 dilution of the homemade 5X extract (~ 93.1 µM) generated a robust response somewhat equivalent to the response elicited by 100 µM eugenol. We, therefore, concluded that most likely eugenol is the sole constituent of Indian clove homemade recipe eliciting repellency activity. Albeit not entirely surprising, it is noteworthy that eugenol is detected by CquiOR73, whereas DEET repellency is mediated by CquiOR136 ^6^.

**Figure 5.**
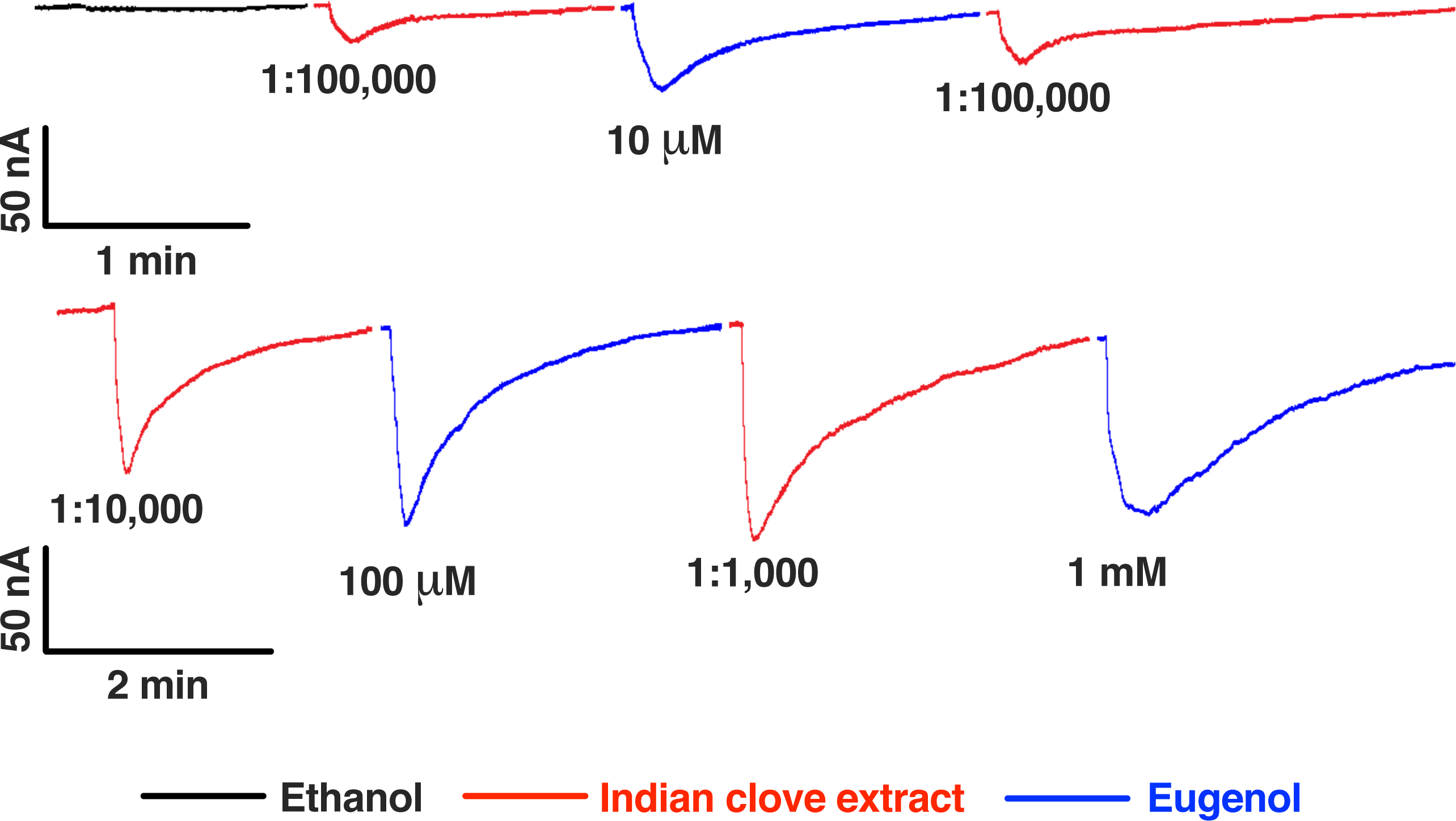
Currents recorded from a CquiOR73/CquiOrco-expressing oocyte challenged with Indian clove extracts and eugenol. 5X Indian clove extracts were diluted with Ringer buffer (1:100,000, 1:10,000, and 1:1,000) and compared to eugenol (10, 100, and 1000 µM = 1 mM). Traces were obtained with different oocytes, but presented here with the same oocyte for comparison.

### *Ae. aegypti* responses to 5X India clove homemade formulation

Lastly, we compared the repellency activity of the 5X homemade recipe with that of 1% DEET against the yellow fever mosquito, which carries dengue, ZIKA, chikungunya, and the yellow fever virus in Brazil. As previously reported, ^17^ protection elicited by 1% DEET is high, but not as high as protection against *Cx. quinquefasciatus*. The homemade extract showed a significantly lower protection that 1% DEET (*n* = 10, P = 0.002, Mann-Whitney test) (Fig. 4C). In conclusion, even a formulation 5X more concentrated than the popular recipe has significantly lower protection than DEET at low dose. It is highly unlikely that the homemade recipe protects the populations from infected mosquitoes.

### Other considerations

It is important to note that older mosquitoes are more likely to transmit diseases than younger ones. If young mosquitoes feed from an infected host, these mosquitoes are more likely to transmit in the next gonotrophic cycle, because there would be enough time for the virus to complete the extrinsic incubation period, ie, the time required for the virus to pass through the midgut barrier into the mosquito’s hemocoel, invade the salivary glands, and replicate to a level that can be infective in the next blood meal ^17^. To fend off these older mosquitoes even higher doses of repellents are required ^17^. For example, it is recommended to use DEET at 20-30% for protection against infected mosquitoes ^17^. In summary, the popular Indian clove recipe is misleading as it does not provide enough protection against mosquito bites. The premise of this recipe is that repellents are expensive, and that the homemade recipe is a cheaper alternative. Our data suggest that even the 5X recipe is ineffective. At the time of this writing the cost of DEET-based products (100 ml, cream with 6.7-7.1% DEET) in Brazil is in average 15.80 BRL (Brazil Real). The cost of 30 g of Indian clove in supermarkets is in average 17.37 BRL, not to mention the other ingredients (alcohol, 3.98 BRL; baby oil, 23.50 BRL). In conclusion, this alternative homemade repellent is a false economy and more importantly a misleading option to prevent mosquito bites.

## Materials and Methods

### Insect Preparations

*Cx. quinquefasciatus* used in this study is derived from a laboratory colony originated from mosquitoes collected in Merced, CA in the 1950s and maintained in the Kearney Agricultural Center, University of California by Dr. Anthony J. Cornel. The “Merced colony” has been maintained in the Davis campus for almost 8 years under a photoperiod of 12:12 (L:D), 27 ± 1°C and 75% relative humidity. The *Ae. aegypti* mosquitoes were from a laboratory colony, commonly referred to as RecLab, which was initiated with mosquitoes collected in a neighborhood (Graças) of Recife, Brazil in 1996. They were kept at the FIOCRUZ-PE facility at 26 ± 1°C, 65-85 % relative humidity, and under a photoperiod of 12:12 h (L:D).

### Chemicals, extractions, and chemical analysis

Hexane (CHROMASOLV®, high performance liquid chromatography grade, CAS# 110-54-3), DEET (PESTANAL®, CAS# 134-62-3), eugenol (CAS# 97-53-0), eugenyl acetate (CAS# 93-28-7), α-humulene (CAS# 6753-98-6), β-caryophyllene (CAS# 87-44-5), and caryophyllene oxide (CAS# 1139-30-6) were acquired from Sigma-Aldrich (Milwaukee, WI); ethanol (high performance liquid chromatography grade, CAS# 64-17-5) was purchased from Fisher Scientific (Hampton, NH). Indian clove buds were acquired from supermarkets in Brazil: Portuense (Product Barcode: 7898290551073), Beija Flor (7896252202360), and Kitano (7891095154081).

Extracts were prepared by weighting at least three pieces of Indian clove buds and then adding enough ethanol to make a final concentration of 0.06 g/ml (original recipe, 1X extracts) or 0.3 g/ml (5X extracts). Vials were closed and kept at room temperature; they were shaken at least twice a day. Upon completing the time of extraction (4 days in the original recipe), the supernatant was separated by filtration and baby oil (Johnson & Johnson, product bar code 7891010877613) was added in the proportion 5:1 (ethanol/baby oil per recipe). For bioassays, the ethanolic extracts were used without further dilution, but for chemical analysis, the extracts were diluted with hexane. For time course analysis, triplicate samples of each of the three brands were prepared, small aliquots were removed at day 4, 5, and 6, diluted 10X with hexane and analyzed immediately.

Gas chromatography (GC) was performed on a 6890 Series GC (Agilent Technologies, Palo Alto, CA) equipped with an HP-5MS capillary column (30 m x 0.25 mm; 0.25 µm, Agilent Technologies). The oven was operated at 70°C for 1 min and increased at a rate of 10°C/min to 290°C, with a final hold of 5 min. The injector and detector were operated at 250°C and helium was used as the carrier gas. The response of the flame ionization detector (FID) was calibrated with authentic samples of eugenol (50-2,000 ng per injection) to generate the following curve: amount of eugenol (ng) injected = 0.06529*Peak Area-27.88 (R^2^ = 0.9920). Gas chromatography-mass spectrometry (GC-MS) was performed on a 5973 Network Mass Selective Detector linked to a 6890 GC Series Plus + (Agilent Technology), which was equipped with the same type of capillary column used in GC analyses. The oven was operated at 70°C for 1 min and increased to 270°C at a rate of 10°C/min. It was held at 270°C for 10 min, with a 10 min post-run at 290°C.

### Measuring behavior

Repellency activity was measured using a previously described surface landing and feeding assay ^6,17,21^. In brief, a mosquito cage frame supported a wood board (30 × 30 × 2.5 cm), which held two Dudley bubbling tubes (painted internally with black glass ink) separated from each other by 17 cm and placed on a transverse plane at the middle line of the wooden board. One side was attached to a mosquito cage (30.5 × 30.5 × 30.5 cm), which housed the test mosquitoes. The side of the wooden board facing the mosquito cage was covered with a red cardstock with openings to allow Dudley tubes to protrude inside the mosquito cage by 5.5 cm. Syringe needles (Sigma-Aldrich, Z108782, 16-gauge) were placed 8 mm above each Dudley tube and inserted 4 cm into the mosquito cage. These syringes deliver CO_2_ (at 50 ml/min) to the cage and held dental cotton rolls placed above the Dudley tubes. Insect pins were placed 1.8 cm above each syringe needle to hold filter paper rings (4 cm × 25 cm, with 1 cm overlapping and stapled together). Water at 38°C was circulated inside the Dudley tubes and once mosquitoes were placed in the test cage, two dental cotton rolls were loaded with defibrinated blood (100 µl, Biological Media Services, UC Davis VetMed shop, catalog #4024), each placed between a syringe and the top of a Dudley tube. Filter paper rings were loaded with 200 µl of solvent only (control) or 200 µl of a test solution (eg, ethanol extract, eugenol, DEET). Solvent was evaporated for 2 min and then the rings were placed inside of the arena being held around each Dudley tube. The procedure was repeated by placing newly prepared filter paper rings and rotating treatment and control so that both sides of the arena were used for control and treatment. Non-blood-fed female mosquitoes (6-7 days old) were separated in aluminum collapsible field cages with green polyester covers (Bioquip, Rancho Cordova, CA) the day before the tests and provided with sugar and water. Assays were recorded with a camcorder equipped with Super NightShot Plus infrared system (Sony Digital Handycam, DCR-DVD 810). The number of mosquitoes that landed on each side of the arena were counted at the end of each assay. To minimize operational errors, filter paper rings were marked with a code of two staples for control and three staples for samples; sample (or solvent) was loaded on the side of the ring with 2-3 staples. The filter paper rings were placed in the arena with the boarder loaded with sample or solvent facing test mosquitoes. For comparison of two repellents (eg DEET vs. extract), the experiments were run in tandem and repeating the following cycles: two replicates with one repellent followed by two replicates with the other repellent. For comparisons, responses were expressed in protection rate, according to WHO and EPA recommendations. Thus, P% = (1-[T/C])X100, where T and C represent the number of mosquitoes landed on the treatment and control sides of the arena, respectively.

### Electrophysiology

The two-electrode voltage-clamp technique (TEVC) was performed as previously described^6,20^. In short, capped cRNA was synthesized by using pGEMHE vectors and the mMESSAGE mMACHINE T7 kit (Ambion-ThermoFisher, Waltham, MA). Purified CquiOR73 cRNA was resuspended in nuclease-free water at 200 ng/µl and 9.2 nl aliquots were microinjected with the same amount of CquiOrco RNA in *Xenopus* laevis oocytes in stage V and VI (Ecocyte Bioscience, Austin, TX). Prepared oocytes were then incubated at 18°C for 3-7 days in modified Barth’s solution (NaCl 88 mM, KCl 1 mM, NaHCO_3_ 2.4 mM, MgSO_4_ 0.82 mM, Ca(NO_3_)_2_ 0.33 mM, CaCl_2_ 0.41 mM, and 4-(2-hydroxyethyl)-1-piperazineethanesulfonic acid, HEPES 10 mM, pH 7.4), which was supplemented with gentamycin 10 mg/ml and streptomycin 10 mg/ml. A stock solution of eugenol (1M) was prepared in dimethyl sulfoxide (DMSO) and then diluted with oocyte Ringer buffer. Ethanolic extracts of Indian clove buds were also diluted with Ringer buffer (NaCl 96 mM, KCl 2 mM, CaCl_2_ 1.8 mM, MgCl_2_ 1mM, HEPES 5 mM, pH 7.6). For TEVC recordings, oocytes were bathed with Ringer buffer at 3.2 ml/min and the holding potential was set at −80 mV. Stimulus (100 µl in 2 s) were injected at 1 cm upstream of the flow. After each stimulus, oocytes were thoroughly washed until a steady baseline was recovered. Chemical-induced currents were amplified with an OC-725C amplifier (Warner Instruments, Hamden, CT), low-pass filter at 50 Hz and digitized at 1 kHz. Data were acquired and analyzed with Digidata 1440A and the software pCLAMP 10 (Molecular Devices, Sunnyvale, CA).

### Graphic preparations and statistical analysis

Graphic illustrations were prepared using Prism 8 for Mac (GraphPad, La Jolla, CA). Original TEVC traces were colored for clarity. A gap between traces indicates that part of the washing phase of the trace was omitted. Statistical analyses were performed with Prism. The specifics for each case are provided in figure legends or directly in the main text. In brief, if a dataset did not pass the Shapiro-Walk normality test, it was analyzed by Wilcoxon matched-pairs signed rank test. One way analysis of variance (ANOVA) was done using Brown-Forsythe and Welch ANOVA tests.

## ACKNOWLEDGMENTS

This work was supported by grants from the National Institutes of Health R01AI095514 and R21AI128931. We thank Fabricio J. Jaciani (FUNDECITRUS, Brazil) and Raquel Silva (University of Brasilia, UNB) for surveying in supermarkets in Araraquara, Sao Paulo and Brasilia the prices of DEET and the ingredients for the Indian clove recipe to combine with the data we collected from Recife, Brazil.

## Author contributions

Designed research: W.S.L. Performed chemical analysis: W.S.L. Performed behavioral experiments: K.T., G.B.F, R.M.R.B., and G.K.B. Molecular cloning and sensory physiology: P.X. Wrote the manuscript: W.S.L. All authors reviewed and approved the final version of the manuscript.

